# Comparing the genomic inbreeding of an isolated rhesus macaque study population to wild populations

**DOI:** 10.64898/2026.06.26.734461

**Authors:** Florence Pautet, Annika Freudiger, Angelina Ruiz-Lambides, Anja Widdig, Harald Ringbauer

## Abstract

Long-term studies of isolated animal populations have greatly improved the understanding of various evolutionary processes. However, potentially elevated inbreeding in those compared to wild populations is a common concern. Conventionally, inbreeding has been investigated using reconstructed pedigrees, but nowadays it can be done directly at the genomic level. Here, we utilise genomic data from an intensively studied isolated rhesus macaque (*Macaca mulatta*) population on the small island Cayo Santiago (Puerto Rico), founded in 1938 with wild animals from India. We quantified inbreeding levels by inferring runs of homozygosity (ROH), i.e., long identical haplotypes inherited from both parents. We identified ROH in 97 ∼5x-coverage genomes from Cayo Santiago and, for comparison, in 79 rhesus macaque genomes from five wild populations from China. Notably, this conventionally considered low-coverage data proved sufficient to infer ROH > 4 centimorgans long after imputing the genomes using a reference panel. Our results revealed that the ROH-derived effective population size on Cayo Santiago, 420 individuals, falls within the ranges we inferred in wild populations. Moreover, a general scarcity of individuals with long ROH in both the Cayo and wild populations indicates very few cases of close-kin breeding, suggesting that mechanisms to avoid close-kin breeding operate in rhesus macaques, in both wild and isolated populations. Taken together, our results suggest that Cayo Santiago remains a representative study population.

## Introduction

Long-term studies of free-ranging animal populations enable researchers to measure natural selection, fitness variation, and life-history trade-offs within the ecological and social environments that drive natural evolutionary processes (reviewed in Sheldon et al., 2022). By providing standardized data over multiple generations and decades, those studies allow scientists to analyse ongoing ecological and evolutionary processes, like aging (e.g., across seven nonhuman primate species: Campos et al., 2022), or variance in fitness (e.g., rhesus macaques: Dubuc et al., 2014; Widdig et al., 2025), as well as the response to environmental change, such as climate change (e.g., meerkats: Paniw et al., 2019 and red deer: Bonnet et al., 2019), or natural catastrophes (e.g., rhesus macaques: Testard et al., 2021).

Long-term studies often focus on isolated populations because they allow precise tracking of animals and facilitate demographic analyses. Prominent examples include populations of Soay sheep on St. Kilda (UK) (Clutton-Brock & Pemberton, 2004) and of red deer on the Isle of Rum (UK) (Pemberton et al., 2022). However, geographically isolated populations may be subject to inbreeding and restricted gene flow, which can cause inbreeding depression (Charlesworth & Willis, 2009). Multiple studies have quantified the degree of inbreeding in isolated study populations and derived individual inbreeding coefficients either from pedigree data (e.g., Walling et al., 2011; Widdig et al. 2017), short-tandem repeats (STRs or microsatellites; e.g., Slate et al., 2000; Widdig et al. 2017), single-nucleotide polymorphism (SNP, e.g., Huisman et al. 2016) or runs of homozygosity (ROH, i.e. genomic regions where identical haplotypes are inherited from each parent due to shared ancestry; e.g., Doekes et al., 2019; Kardos et al., 2018; Stoffel et al., 2021). These studies found that inbred animals had lower survival rates and lower lifetime breeding success (e.g., red deer: Huisman et al., 2016; Slate et al., 2000; and Soay sheep: Stoffel et al., 2021). Therefore, understanding the prevalence of inbreeding in isolated study populations is of high interest for informing population management decisions and validating the representativeness of the population for a given species.

Runs of homozygosity (ROH) are a particularly powerful signal to study various aspects of inbreeding. Because recombination continuously breaks up stretches of genomic material, long ROH within an individual genome indicate close kinship between the parents. Shorter ROH reflect more distant coalescences (Kardos et al., 2016) and therefore yield insights into recent demographic events (Ceballos et al., 2018). They can be used to estimate effective population sizes (Ringbauer et al., 2021), which reflect the number of individuals effectively contributing genes to future generations. The effective population size is often substantially smaller than the census population size (Frankham, 1995; Palstra & Ruzzante, 2008), although the relationship between effective and census sizes remains complex (Palstra & Fraser, 2012). Several bioinformatics methods can identify ROH, including sliding-window-based approaches (PLINK: Chang et al., 2015; Purcell et al., 2007), Hidden Markov Chains (bcftools/ROH: Narasimhan et al., 2016), and linkage information from reference panels (hapROH: Ringbauer et al., 2021). Detecting ROH can be challenging, particularly when genomic resources are suboptimal (e.g., a fragmented reference genome) or genomic coverage is low. But increasingly available genomic resources make identifying ROH a routine task in modern (reviewed in Ceballos et al., 2018) and ancient humans (Ringbauer et al., 2021), livestock (reviewed in Peripolli et al., 2017), and, increasingly, non-model species (Brüniche-Olsen et al., 2018). These genomic applications have yielded insights into inbreeding depression (Doekes et al., 2019; Stoffel et al., 2021), demographic history (Ceballos et al., 2019; Hewett et al., 2023), and conservation management (Mueller et al., 2022; Robinson et al., 2019).

The rhesus macaque (*Macaca mulatta*) population on the island of Cayo Santiago, off the coast of Puerto Rico, is a prominent example of an isolated long-term study population. Originally descended from 409 animals, collected in 1938 at different locations in India (Rawlins & Kessler, 1986), this free-ranging island population combines the ecology of a natural setting with the demographic depth of a long-term pedigree record. The population has become an intensively studied model for understanding how ecological pressures and social structure shape evolutionary processes in primates. Various studies on the Cayo Santiago macaques have focused on questions such as the evolution of social bonds (Clive et al., 2023; Ellis et al., 2019; Testard et al., 2021), life-history trade-offs (Patterson et al., 2024), ageing (Testard et al., 2024; Watowich et al., 2024), and the genetic architecture of complex traits (Costa et al., 2025). Despite a relatively large long-term census size, the population has remained genetically isolated since its foundation (Widdig et al., 2017). Therefore, founder effects and genetic drift, together with breeding among close kin, may have greatly reduced its genetic variability, raising questions about whether the population on Cayo Santiago is genetically comparable to wild populations.

Prior studies have started to address the question of inbreeding and genetic diversity in the Cayo Santiago population, as this could have important implications for its status as a model system. Genetic markers inferred from red blood cell phenotypes documented substantial genetic diversity in 1972 (Duggleby, 1978a). Genetic variation likely declined due to substantial animal removals in 1972 (Duggleby, 1978b; reviewed in Widdig, Kessler, et al., 2016). Nevertheless, subsequent population genetic assessments based on STR markers revealed sufficient genetic variation and low rates of inbreeding (Kanthaswamy et al., 2017; Widdig et al., 2017). Specifically, Kanthaswamy et al. (2017) reported an observed heterozygosity of 0.72 and that social groups did not differ genetically from each other, likely due to gene flow through dispersing males. Furthermore, Widdig et al. (2017) reported evidence of limited close-kin inbreeding, attesting to inbreeding loops in 21 of 2,669 analysed individuals (0.8%) using complete three-generation pedigrees, and in 45 of 609 individuals (7.4%) using complete four-generation pedigrees. Most close inbreedings involved individuals related via the paternal kin line, particularly matings between paternal half-siblings (18 of 21 cases). Overall, inbreeding levels did not change significantly over the 23-year study period. Moreover, Widdig et al. (2017) found evidence of inbreeding avoidance, as the observed mean relatedness of reproducing pairs was significantly lower than expected, considering the degree of male reproductive skew, extra-group paternity, and natal breeding. Finally, there was no significant association between inbreeding and early infant death or survival until maturation. However, genome-wide data might more precisely quantify inbreeding than partial pedigrees or microsatellites (Huisman et al., 2016; Kardos et al., 2015) and elucidate patterns of general background relatedness.

To explore the extent of inbreeding on Cayo Santiago with greater resolution, we reanalyse published genomic data from this population (Freudiger et al., 2025) and compare them with published genomes from wild rhesus macaques collected across China (Liu et al., 2018). Due to the absence of genomic data from wild Indian rhesus macaques (as of 2026), the Liu et al. (2018) study is the closest available genomic comparison to the Indian-derived population of Cayo Santiago. The Chinese and the Indian rhesus macaque lineages were estimated to have split around 160k years ago (Hernandez et al., 2007) and subsequently diverged due to geographic isolation. However, Chinese- and Indian-derived rhesus macaques are classified as the same species (Champoux et al., 1997), as they remain capable of interbreeding and share morphological features (Hamada et al., 2005). In their study, Liu et al. (2018) divided their Chinese sample set into five subspecies. Although the formal classification of *Macaca mulatta* remains debated due to the limited genetic differentiation among its subspecies (e.g., Terbot et al. 2025), we retain the subspecies labels here to distinguish geographically separated populations. However, we stress that we use them only as labels for Chinese populations, referring to geographic ranges and without assigning any taxonomic significance.

Our study aims to compare patterns of long and short ROH in the genetically isolated Cayo Santiago population with those of published wild macaque populations, to provide insight into relative patterns of both close-kin breeding and recent effective population sizes. Assessing these patterns allows us to compare the Cayo Santiago population with various wild animal populations and, at least from a genomic perspective, to determine whether an isolated study population such as Cayo Santiago can remain representative of wild populations.

## Data & Methods

### Study species

Rhesus macaques (*Macaca mulatta*) are native to Asia and have the largest geographic range of any nonhuman primate, extending from India to China and south into Vietnam and Thailand (Xue et al., 2016). They live in large groups comprising numerous males and females, along with their offspring (Widdig, Langos, et al., 2016). Females are philopatric, i.e., they remain in their birth group throughout their lives and form matrilineal kin groups, whereas males disperse from their birth group around puberty to breed elsewhere (Weiß et al., 2016). This sex-biased dispersal has been discussed as one of the main mechanisms of inbreeding avoidance (Pusey & Wolf, 1996), as kin availability is similar for both sexes at the time of maturation and before male dispersal (Widdig, Langos, et al., 2016). Rhesus macaques are seasonal breeders and mate with multiple sexual partners (Hoffman et al., 2008). Male reproduction is skewed: some males produce many offspring, while most produce none. Such skew has been observed in the annual reproduction rate in long-term data and from a lifetime perspective (Dubuc et al., 2014; Widdig et al., 2004, 2025). As a consequence, male reproductive skew will affect kin structure and sociality within social groups (Widdig, 2013).

### Study populations

Our focal interest lies in the rhesus macaque population on Cayo Santiago, a 15.2-ha island off the southeast coast of Puerto Rico, USA. It was founded in 1938 by introducing 409 wild-born Indian-origin rhesus macaques collected from seven different provinces in central India (Rawlins & Kessler, 1986). Since then, no new individuals have been introduced. The population is managed by the Caribbean Primate Research Center (CPRC) and maintained in semi-natural conditions, i.e., animals are provided with a daily supply of commercial monkey diet, but still spend about 20-50% of their feeding time foraging on natural vegetation (Kuthyar et al., 2022; Marriott et al., 1989). Individuals are identified, genetically sampled, and continuously monitored across their lifespans, allowing researchers to directly link variation in genotype, phenotype, behaviour, and fitness (reviewed in Widdig, Kessler, et al., 2016). To maintain the population at reasonable levels, individuals and, occasionally, social groups, are regularly removed as the primary demographic management strategy (Hernandez-Pacheco et al., 2016). Most notably, multiple large culls reduced the population from 800 in 1968 to around 300 individuals in 1972 (Duggleby et al., 1986), and 596 individuals (around 50% of the population) were removed at once in 1984 (Hernandez-Pacheco et al., 2016). In the present study, we used animals born between 1981 and 2009.

To provide context from rhesus macaques in natural habitats, we analysed 79 wild rhesus macaques from 17 local wildlife rescue centres in China (**Fig. S1**) using genomic data from a previous study (Liu et al., 2018). Liu et al. (2018) divided their samples into five subspecies. While this classification is not generally accepted (Terbot et al. 2025), we retain the original naming convention used by Liu et al. (2018) to distinguish between geographically separated populations.

### Data preparation and bioinformatic processing

For the Cayo Santiago population, we used genomic data produced by Freudiger et al. (2025). Sequencing methods, genotype imputation, and variant calling are described in the original study. In summary, they selected a subset of the pedigree by choosing one individual and adding all of that individual’s 1^st^-degree relatives to the sample. In the next two rounds, they added all 1^st^-degree relatives of the individuals identified in the previous round. Samples were sequenced to a median genomic depth of 5.06× (range: 3.05-37.94×). After standard quality filtering, the sequences were aligned to *Mmul10* (Warren et al., 2020), then imputed and phased using a reference panel of 741 Indian-origin *M. mulatta* individuals from mGAP 2.4 (Bimber et al., 2019). Of 102 Cayo genomes, 98 passed quality tests (Freudiger et al., 2025). Additionally, we excluded one individual (194298) after ROH calling due to signs of DNA contamination. The final dataset contained 97 individuals and 7,007,674 biallelic SNPs.

For the Chinese rhesus macaques, we used a published VCF file containing previously called variants (Liu et al., 2018). As described in the original study, these samples derive from both blood and muscle tissue and were sequenced to high coverage (> 20×) for 9 genomes and to 8-12× coverage for the remainder. After quality filtering, Liu et al. (2018) aligned the reads to *Mmul8* (Zimin et al., 2014) before variant calling. To standardise both datasets, we removed multi-allelic SNPs and indels, retaining only genotypes with quality scores ≥ 20. We then lifted the genomes over from *Mmul8* to *Mmul10* using *LiftoverVcf* from *GATK* (McKenna et al., 2010), retaining only SNPs mapped to one of the 20 autosomal chromosomes. This processing resulted in a dataset comprising 79 individuals and 20,043,686 biallelic SNPs.

### Calling runs of homozygosity (ROH)

According to Silva et al. (2024), *BCFtools/ROH* and *PLINK* perform similarly, although *BCFtools* is more likely to miss short ROH, while *PLINK* is more prone to overestimate long ROH. We chose *BCFtools/ROH* v1.22 (Narasimhan et al., 2016) because it allowed us to use both the genotype probabilities and the linkage-disequilibrium-based genetic map (Freudiger et al., 2025), and was less likely to overestimate close-kin breeding. To account for recombination hotspots, inferred ROH segments were measured in centimorgans (cM). Because occasional genotyping errors can split long ROH segments into mosaics of shorter inferred ROH (Ringbauer et al., 2021), we merged nearby ROH segments (each > 0.5 cM long and separated by a gap < 1 cM). For downstream analysis, we retained segments > 4 cM, as shorter segments are challenging to detect and prone to false positives (Ringbauer et al., 2021). We manually inspected 30 genomes, including the 10 most and 10 least inbred, by comparing the inferred ROH with the heterozygous SNP density across the genome. We found that ROH were readily visible in these plots (see an example in **Fig. S2**), allowing us to verify the inferred ROH. Moreover, we could assert that no ROH were false positives due to low SNP density and checked the individuals without detected ROH for false negatives. Overall, this visual inspection revealed excellent performance of our ROH calling pipeline for ROH > 4 cM. Notably, this manual verification led us to exclude one genome (194298) with a contamination signature (**Fig. S2b**).

### Classifying ROH

Following previous approaches (Racimo et al., 2020; Ringbauer et al., 2021), we grouped inferred ROH segments for each individual into bins by genomic length (here: 4-8, 8-12, 12-16, 16-20, > 20 cM) and summed the overall length of all ROH within each bin. To quantitatively assess the timescale of coalescence events underlying ROH, we calculated the densities of coalescence times for each bin, using the genomic map of rhesus macaques (**Fig. S3**). We assumed a constant size N_e_ = 400, but we note that the exact value does not substantially change qualitative patterns. For example, 95% of ROH > 20 cM originate from within 12 generations, while 50% of 4-8 cM long ROH originate between 9 and 30 generations ago. These time intervals are broad; thus, we stress that it is impossible to map an ROH of a given length to a single time point. Nevertheless, the rapid recombination clock provides strong upper bounds on the signal’s age (**Fig. S3**; Ceballos et al., 2018).

### Simulating ROH distributions

To verify the theoretical framework and explore stochastic variation around the analytically accessible averages, we simulated ROH for various scenarios of parental relatedness. Specifically, we used i) *pedsim* (Caballero et al., 2019) to simulate ROH for offspring of various degrees of close-kin breeding (full- and half-siblings; avunculars; 1st-, 2nd- and 3rd-cousins) and ii) *msprime* (Baumdicker et al., 2022) to simulate ROH for various effective population sizes, assuming a randomly mating population with a constant size N_e_ in [100, 200, 500, 1000, 2000]. The resulting ROH distributions (**Fig. 1c,d**) confirmed the theoretical expectation that the length bins contain information about different time depths of inbreeding loops.

**Figure 1:**
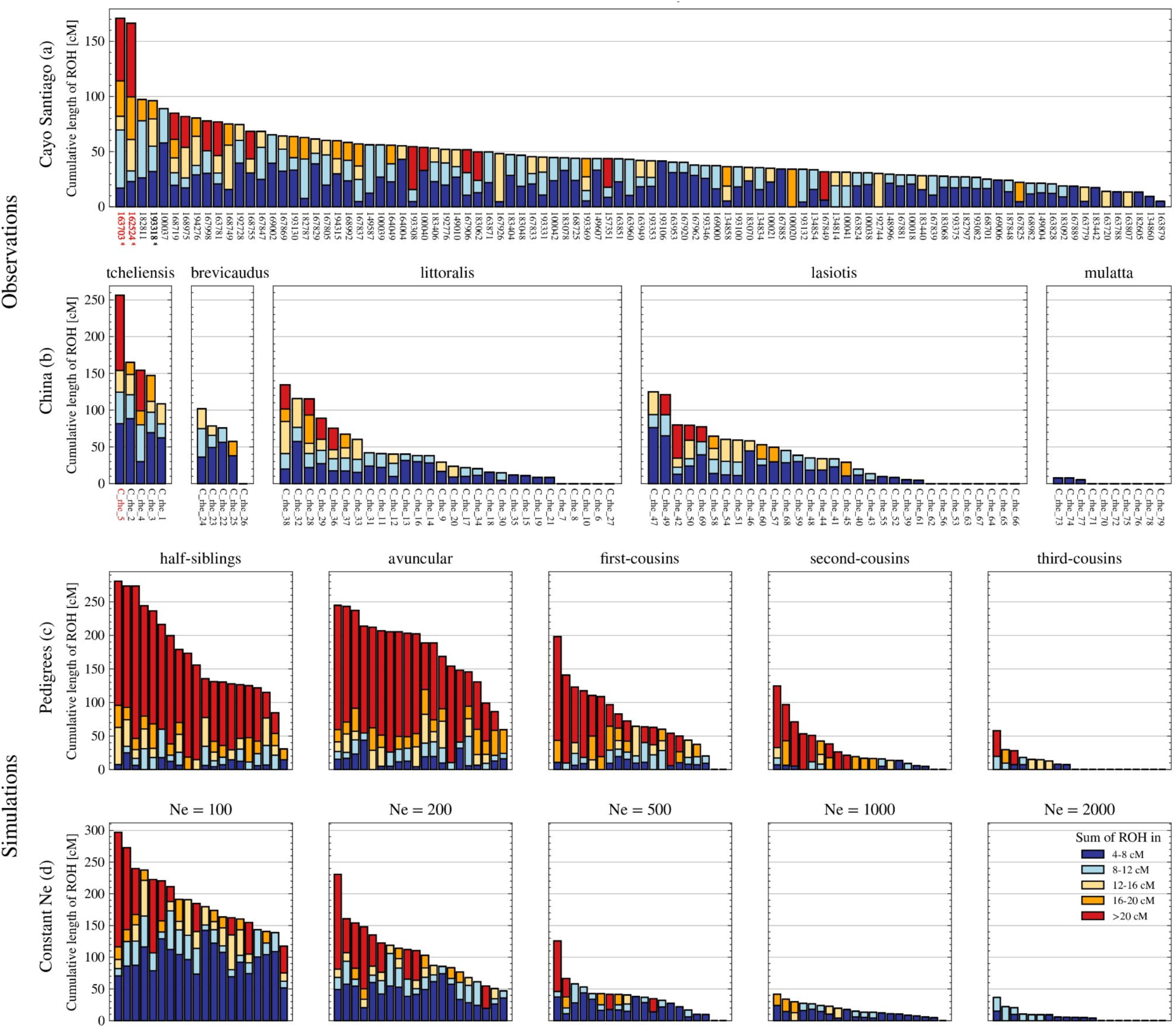
ROH summary for both observed and simulated individuals. Each vertical bar represents one individual (labelled on the x-axis). The colored bars represent the summed ROH length (in cM) within five bins (see legend). Red segments correspond to the longest ROH category (> 20 cM), indicative of close inbreeding, while blue bars correspond to short ROH (4-8cM), indicative of more distant parental relatedness. Note that different panels have different y-axis scales. **(a)** and **(b):** Inferred ROH in the samples from Cayo Santiago and from five wild Chinese populations. Individuals with > 5% of their genome covered by ROH segments > 16 cM (implying they are likely descendants of close-kin unions) are marked using red labels. Known half-cousin offspring (according to the pedigree) are marked using a bold label and an asterisk. (**c):** Simulated individual ROH for various degrees of parental relatedness. **(d):** Simulated ROH within a randomly mating diploid population of constant effective population sizes Ne.

### ROH- and pedigree-based estimates of inbreeding

To quantify inbreeding, we used the genomic inbreeding coefficient *F_roh*, defined as the proportion of an individual’s genome that is autozygous (McQuillan et al. 2008). For each individual, we calculated *F_roh* by dividing the sum of the lengths of its ROH segments longer than a given cutoff by the total genome map length. We explored various cutoffs (4, 8, 12, 16 cM) to assess their impact on the estimate and denote the resulting coefficient by *F_rohₓ*, where x refers to the cutoff value.

The measure *F_roh* is expected to correlate with the pedigree-based estimate *F_ped* introduced by Wright (1922) and Malécot (1948). Following a previous approach (Freudiger et al., 2025), we used the program *TRACE* v0.1.0 (Westphal et al., 2023) to calculate pedigree-based relatedness estimates by summing over all links in the pedigree for 12,049 members of the Cayo Santiago rhesus macaque population, including 11,805 known mother-offspring dyads and 4,986 known father-offspring dyads. We applied *TRACE* to the 63 individuals in our genomic dataset for whom we had at least a complete three-generation pedigree and calculated their inbreeding coefficient (*F_ped*) as half the pedigree-relatedness coefficient (*r_ped*) between their parents. We then categorised those individuals into distinct kin classes based on their pedigree. 30 of 63 individuals were inbred via multiple (mostly distant) loops; in these cases, we classified them based on the most recent inbreeding (i.e., the one that contributed the most to the overall inbreeding coefficient).

### Estimating effective population size

We inferred the effective population size using a maximum-likelihood estimation (MLE) framework, using ROH segments 4-12 cM long as input. The likelihood function derives from an established formula (Browning & Browning, 2015; Ringbauer et al., 2017) that predicts the expected numbers of autozygous segments of various lengths under the assumption of a randomly mating population of constant size N_e_. The inferred N_e_ is the one that maximises the likelihood of the observed ROH lengths across all individuals. Confidence intervals of the estimates were obtained using Wilk’s theorem and correspond to the range of values within χ^2^_1, 0.95_ /2 = 1. 92 units of the maximum-likelihood estimate.

## Results

### Incidence of close-kin breeding measured by long ROH

According to the pedigree, none of our study animals from Cayo Santiago has parents related within three degrees of kinship; the most inbred individuals in our sample are three offspring from half-cousin matings (fourth-degree relatives). This observation is corroborated by the inferred long ROH (defined as > 16 cM): 70 out of 97 animals (72.2%) lack any long ROH segments (**Fig. 1a**). Only two individuals stand out, with *F_roh₄*=11% for both and *F_roh₁₆*=6.0% and 7.1%, respectively. They correspond to two of the three known half-cousin offspring. The third half-cousin offspring still ranks at position 4 with *F_roh₄*=6.5% and *F_roh₁₆*=1.1%. The variance within parental kin class is entirely expected due to random recombination, as seen in the simulated ROH of close-kin offspring (**Fig. 1c** and **Fig. S4**). The inferred ROH confirms that the potentially incomplete pedigree does not miss any cryptic close-kin breeding in our sample.

We observed a similar pattern of long ROH (> 16 cM) in the Chinese samples, in which only one of 79 individuals (1.3%) has > 5% of their genome covered by long ROH (**Fig. 1b**). While we do not have pedigree information for the Chinese samples, its inbreeding coefficients *F_roh₄*=17% and *F_roh₁₆*=6.8% suggest a second-degree parental relationship (e.g., half-siblings).

### Background relatedness measured by shorter ROH

The patterns of shorter ROH indicate a rather consistent level of background relatedness on Cayo Santiago: 94% of the individuals have at least one ROH segment between 4-8 cM, and 73% have at least one segment between 8-12 cM. Overall, the number of short ROH segments does not vary substantially across individuals (**Fig. 1a**). This pattern is consistent with the reconstructed pedigree, which indicates that 70% of the individuals considered have parents related within the last 5 generations (10th degree of kinship) (**Tab. S1**).

In the Chinese samples, we find that patterns of background relatedness vary markedly across populations, with individuals within each population showing mostly consistent levels of short ROH. *M. m. littoralis* and *M. m. lasiotis* present a similar ROH profile to the Cayo Santiago population, whereas we hardly detect any ROH in the wild Chinese *M. m. mulatta*. However, while we find at least one ROH segment ≥4 cM in each individual from Cayo Santiago and in most of the simulated ones (99% for Ne=500 and 90% for Ne=1000), this is not the case in the wild populations. In both populations with the closest ROH profiles to Cayo Santiago, *M. m. littoralis* and *M. m. lasiotis*, we observe, respectively, 5/28 (17%) and 8/31 (25%) individuals without any ROH segments. We even find one such animal among the five from *M. m brevicaudus*, where the average ROH amount is otherwise twice as high. One plausible explanation for sporadic individuals without ROH > 4 cM is occasional mating with long-distance migrants, resulting in outbred offspring with little ROH.

### Comparison of *F_ped* and *F_roh*

Both *F_roh* and *F_ped* quantify inbreeding at the individual level. On Cayo Santiago, using the least stringent cutoff of > 4 cM, we find that *F_roh₄* (mean=3.40%, median=2.96%) is consistently higher than *F_ped* (mean=0.35%, median=0.10%; **Fig. 3** and **Tab. S2**). This bias is not unexpected and has two potential explanations. First, *F_ped* does not account for background relatedness. In a small population or after a founder event, even unrelated individuals share some DNA co-inherited from distant common ancestors. Second, the pedigree contains gaps. While the mother of an individual identified via observation has been confirmed genetically in the majority of cases (98.7%; Widdig et al. 2017), the sire cannot be inferred from observations. Systematic genetic paternity testing has been conducted on the island since 1992, but it does not apply to earlier generations in the Cayo Santiago pedigree, and some cases remain unsolved despite these efforts. Consequently, for the 63 animals with complete three-generation pedigrees, on average, half of the great-grandfathers are unknown.

Using a longer ROH length cutoff for *F_roh* yields a higher correlation with *F_ped*; we observed the highest Pearson correlations for ROH > 8 cM (r=0.69) and > 16 cM (r=0.70), compared to > 4 cM (r=0.66) (**Tab. S2**).

### Effective population size estimates

The inferred prevalence of ROH across different lengths on Cayo Santiago falls within the range we observed in various Chinese populations (**Fig. 2**). Consequently, the same accounts for the effective population size (**Tab. 1**). We estimate N_e_ = 421 (95% CI: 384-464) for Cayo Santiago, and sizes ranging from 121 to 536 for four of the five wild populations. Only *M. m. mulatta* stands out, with an estimated size N_e_ = 6,862 (95% CI: 2,638 - 27,623), which is an order of magnitude larger than that of the other populations. We stress that the statistical certainty of those estimates depends on the total amount of ROH, i.e., on the number of samples times the average amount of ROH per sample. Therefore, for the same number of samples, a population with a large effective population size, such as *M. m. mulatta*, will have significantly larger confidence intervals than a small population such as *M. m. tcheliensis*.

**Figure 2:**
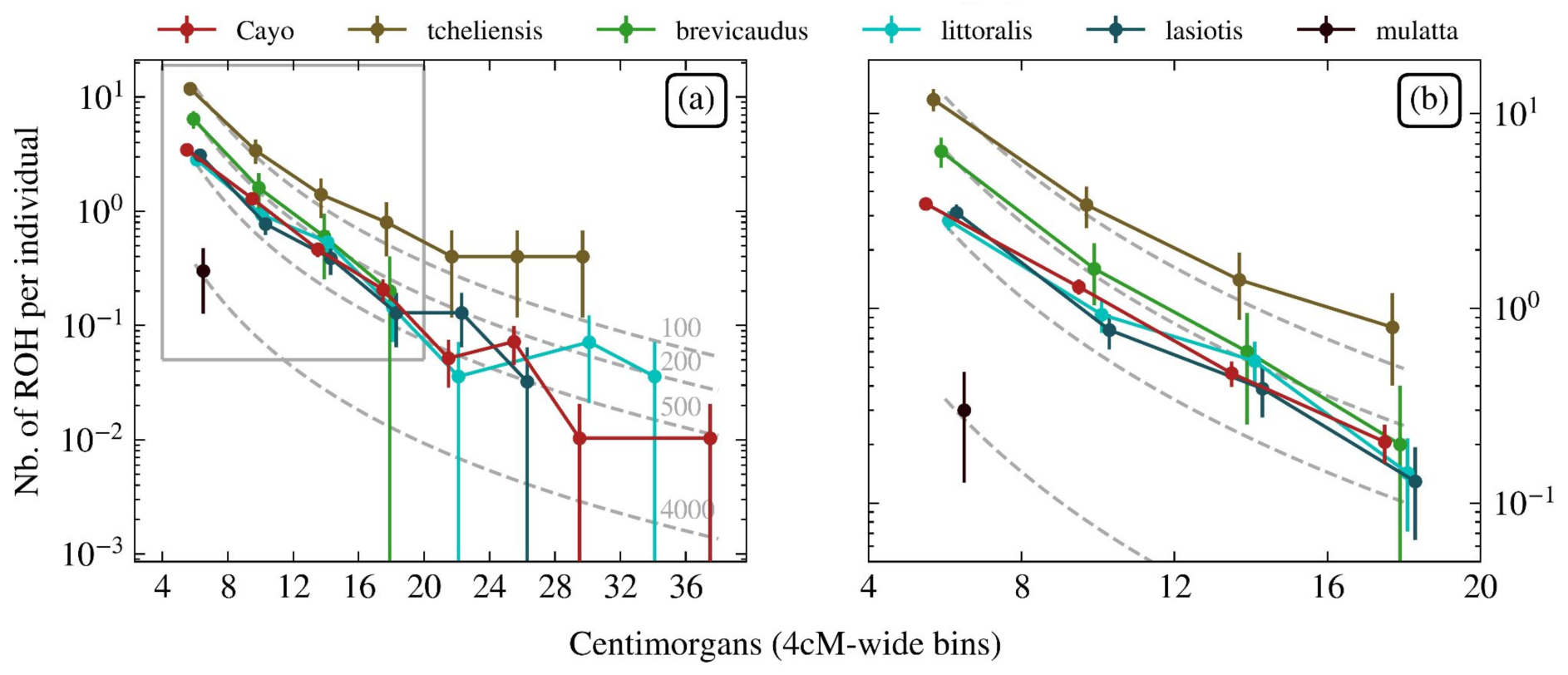
Average rate of ROH segments per population across various lengths. **(a)** We depict the rates of ROH segments per population in 4cM-wide bins. Confidence intervals correspond to 68% coverage intervals assuming a Poisson distribution in each bin. Point estimates (circles) are placed at the centre of each bin, with jitter added to the x-coordinate for visual separation. Dashed grey lines indicate the theoretical distribution for panmictic populations of various constant effective sizes. Due to the logarithmic y-scale, bins without ROH do not appear as points. The grey frame depicts the area expanded in panel (**b)**. **(b)** Zoom of the shorter ROH lengths only. They are more numerous and therefore have relatively higher sampling certainty.

**Figure 3:**
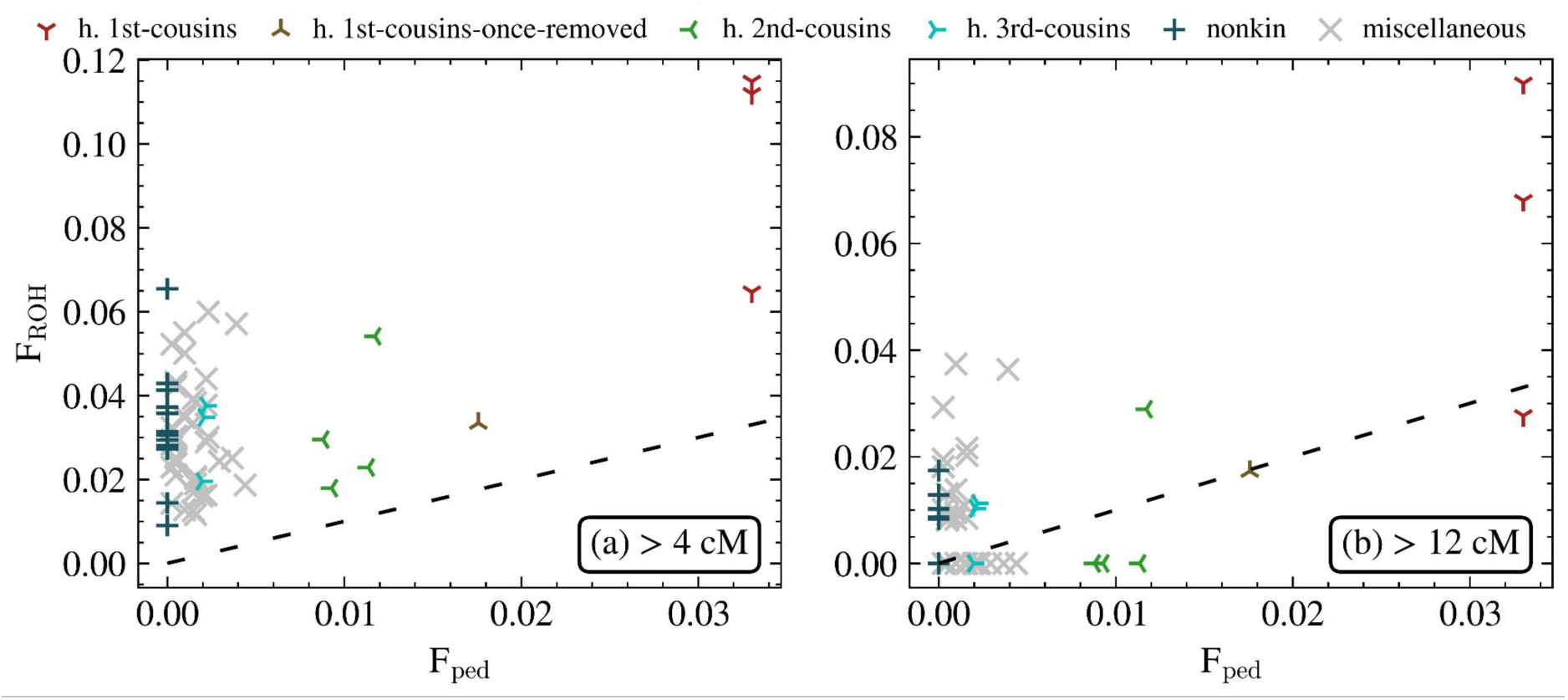
Comparison of inbreeding coefficients F_ped and F_roh on Cayo Santiago. Each dot represents an individual from Cayo Santiago with a complete 3-generation pedigree. The labels refer to the degree of parental relatedness based on the pedigree. Only the closest kin classes are labelled; all more distant relationships fall under “miscellaneous” (see **Tab. S1** for more details). *F_roh* was computed using all ROH longer than **(a)** > 4 cM and **(b)** > 12 cM. The dashed line depicts the agreement line *F_ped* = *F_roh*.

**Table 1:**
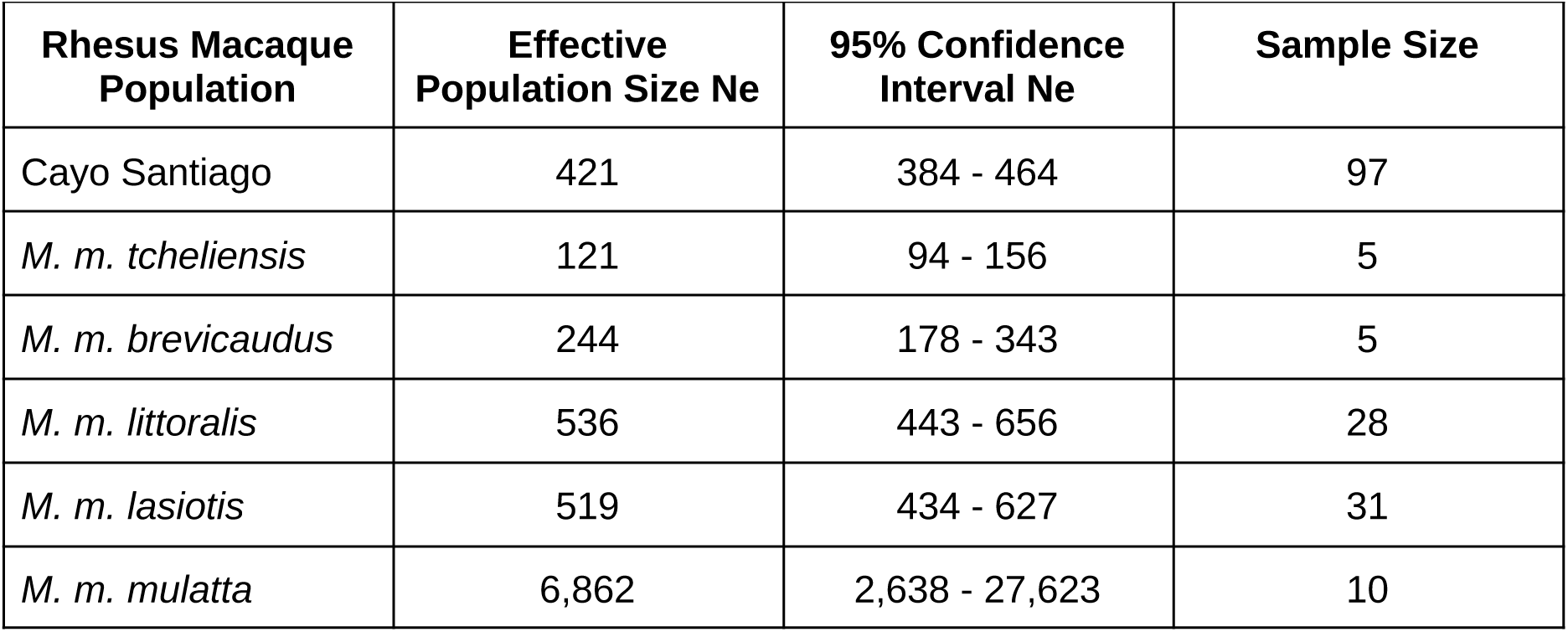
Estimated effective population sizes for the different studied populations. These estimates are based on fitting inferred ROH 4-12cM long with a likelihood framework (see **Methods**). The column Sample Size denotes the number of individuals whose inferred ROH we used to estimate Ne.

## Discussion

In this study, we investigated inbreeding patterns in a long-term-isolated study population by inferring ROH in 97 rhesus macaques from Cayo Santiago and comparing them with those of 79 individuals from five wild Chinese rhesus macaque populations. We used ROH to investigate two aspects of inbreeding: first, long ROH segments that signpost close-kin breeding; second, short ROH segments associated with background relatedness and effective population size. Remarkably, we find that the Cayo Santiago population falls within the range of wild populations in both respects, revealing that the genetically isolated population of Cayo Santiago is not more inbred than several wild rhesus macaque populations in China.

First, given the low frequency of long ROH (> 16 cM), we find that close-kin breeding is rare in both the Cayo Santiago and wild rhesus macaque populations. Across all 97 individuals from Cayo Santiago, we identified only three cases of possibly fourth-degree relatedness among parents (e.g., half-cousins), and no closer-kin matings. We observed a similar lack of close-kin breeding, even in the wild population with the smallest effective population size (*M. m. tcheliensis*, inferred N_e_ ∼120). This general lack of close inbreeding in Cayo Santiago and wild rhesus macaque populations is consistent with prior studies (Widdig et al. 2017). Our results suggest that mechanisms of inbreeding avoidance are also effective in small or isolated rhesus macaque populations. There are various potential mechanisms underlying this lack of inbreeding: Various species have different *pre-copulatory* mechanisms of inbreeding avoidance, mainly through (sex-biased) dispersal (e.g., black bears: Costello et al., 2008; prairie-dogs: Dobson et al., 1997 and great tits: Szulkin & Sheldon, 2008) and kin discrimination in mating partner choice (e.g., storm petrels: Bonadonna & Sanz-Aguilar, 2012) or a combination of multiple mechanisms (e.g., elephants: Archie et al., 2007; baboons: Galezo et al., 2022 and mountain gorillas: Morrison et al., 2023). Inbreeding can also be prevented after copulation; for example, a recent baboon study demonstrated that *post-copulatory* differences in gene expression and vaginal tract pH in mated females reduce the probability of conception by sperm from genetically similar males (Petersen et al., 2026). Similarly, horses have a lower conception rate if females are exposed to odour cues of genetically similar males during their conceptive phase (Burger et al., 2017). Generally, a previous meta-analysis suggested that avoidance through mate choice evolves as a mechanism to prevent inbreeding depression, particularly when related sexual partners regularly encounter each other and inbreeding costs are high (Pike et al., 2021). Rhesus macaques, specifically, were shown to discriminate unfamiliar paternal kin from nonkin solely based on single phenotypic cues, namely facial images (Pfefferle, Kazem, et al., 2014) and calls (Pfefferle, Ruiz-Lambides, et al., 2014), an ability which could facilitate inbreeding avoidance.

Second, shorter ROH (4-12cM) provide information about the background relatedness, i.e., the baseline of genetic similarity between two distinct individuals from a population. In this respect, we find the patterns of shorter ROH to be remarkably consistent within each population (**Fig. 1**), indicating distinct, persistent levels of background relatedness. The prevalence of short ROH in the Cayo Santiago population (**Fig. 2**) and its effective size, which we estimated at N_e_∼420 fall within the range of variation we observe in wild populations in China (**Tab. 1**). Apart from the Chinese population *M. m. mulatta,* which has a significantly larger estimated population size (N_e_∼6,800), all other populations have N_e_ between ∼120 and ∼520. Notably, unlike on Cayo Santiago and in our simulations (which assume randomly mating populations of fixed sizes; **Fig. 1d**), we observe several animals from wild Chinese populations lacking any ROH ≥ 4 cM. Those evidently outbred animals could suggest gene flow from long-distance migration among the Chinese populations. Although male dispersal also exists on Cayo Santiago (Widdig et al., 2017), it is limited to a few groups that have not differentiated significantly.

Our estimated effective population sizes can be compared with prior knowledge of the populations. Unsurprisingly, the Chinese population with the highest ROH prevalence and, consequently, the smallest effective size also occupies the smallest territory (**Fig. S1**) and suffers from habitat fragmentation and contraction (Lu et al., 2007). For the Cayo Santiago population, census data report that the total number of adults (defined as > 4 years old) globally increased across the last seven decades, rising from 180 to 1,300 individuals, with an arithmetic mean of 530 (Cooper et al., 2024). The reproductive skew in this species, especially among males (Dubuc et al., 2014; Widdig et al., 2025), tends to reduce the effective size compared to the census size (Miller et al., 2009), and multiple generations alive are not captured by the single-generation effective size estimate (Ruzzante et al., 2016). Thus, the recent demographic population size estimates match our ROH-based effective population size estimate of N_e_∼420. Finally, in an expanding population, we would expect more distant coalescence events, resulting in an excess of short ROH compared to a model population of constant size. Yet a constant-population-size model accurately captures the observed ROH distribution (**Fig. 2**). This pattern suggests that additional demographic factors beyond overall population size (e.g., subdivision) are at play. Moreover, the ROH-based estimates of effective population size N_e_ for Cayo are similar to prior N_e_ estimates based on IBD segments shared among individuals (Freudiger et al., 2025). These IBD-based estimates yielded a N_e_ range of 233-425 (95% CI), overlapping with the ROH-based estimate of 421. This congruence is reassuring, as IBD and ROH segments are closely related concepts: both are long shared haplotypes (IBD between two individuals and ROH within one individual) that reflect recent co-ancestry patterns, and ROH segments directly result from IBD segments shared between the parents.

The inbreeding patterns we document in rhesus macaques align with prior inbreeding studies in natural populations, both open and isolated, which reported that inbreeding levels do not necessarily depend on a population’s genetic or geographic isolation. For example, while some studies found some level of inbreeding in captivity (e.g., Fredrickson & Hedrick, 2002) and isolated populations (e.g., Marshall & Spalton, 2000), others detected negligible inbreeding in similarly sized populations (e.g., Overall et al., 2005; Rzewuska et al., 2005). However, high or moderate inbreeding has also been observed in wild populations (e.g., Kardos et al., 2018; Prado-Martinez et al., 2013), often linked to reduced fitness (e.g., Amos et al., 2001; Rijks et al., 2008). Together, these studies and ours highlight that isolated populations are not necessarily more inbred than open populations, reinforcing that isolated populations can, in principle, be a valuable study system for evolutionary research.

Our study highlights the general utility of ROH for examining various aspects of inbreeding in both open and isolated populations. Importantly, we show that inferring ROH enables comparisons of inbreeding patterns between long-term field studies and less-documented wild populations, even when pedigree is lacking. On Cayo Santiago, we observed *that F_roh* was systematically higher than *F_ped*, primarily because the pedigree lacks distant parental relationships. Further, we note that the correlation between *F_ped* and *F_roh*, as well as the interpretation of *F_roh*, depends on the length cutoff used to compute *F_roh*. Shorter ROH capture older coalescent events not necessarily recorded in the pedigree. In contrast, longer ROH reflect only recent inbreeding loops, which might be of greater interest when studying inbreeding depression or close-kin breeding (Kyriazis et al., 2025). Our study also provides a successful blueprint for inferring ROH from genomic data in rhesus macaques, as our pipeline can be readily applied to other rhesus macaque populations and, with some adjustments, to other species. Importantly, we demonstrate the utility of genome imputation for inferring ROH, even at ∼5× coverage, which is not conventionally considered high for this purpose (Silva et al., 2024). We caution, though, that quality control of inferred ROH is essential for each lower-coverage application.

There are several limitations to our analysis. First, one might question the representativeness of the sample, given the sampling strategy employed by Freudiger et al. (2025), which focused on a subset of the Cayo Santiago pedigree. However, there is no straightforward reason why this subset should have exceptional inbreeding patterns. Second, we used a linkage disequilibrium-based genetic map (Freudiger et al., 2025) with lower resolution than human recombination maps, and linkage disequilibrium-based methods such as *LDhelmet* (Chan et al., 2012) may exhibit biases (Dapper & Payseur, 2018; Raynaud et al., 2023). Third, although broad-scale recombination rates are conserved between Chinese- and Indian-origin macaques, there are notable differences in the fine-scale recombination landscapes of Chinese- and Indian-derived rhesus macaques (Terbot et al., 2025). An overestimated linkage disequilibrium would result in shorter genetic lengths and thus an underestimation of inbreeding. However, errors at fine genomic scales should average out when considering multiple centimorgans and are very unlikely to affect our analysis. Fourth and probably most important, we compare Chinese and Indian-origin rhesus macaque populations. While they belong to the same species and exhibit modest genetic differentiation (F_ST_ ∼0.15; Heenkenda et al., 2026), there may be meaningful differences in inbreeding patterns among wild Indian populations that we could not observe here due to the lack of publicly available genomic reference data. We hope that future genomic resources will help to close this gap.

Our findings provide directions for future studies. First, the lack of close-kin breeding we documented suggests, in line with prior evidence, that rhesus macaque populations have mechanisms to avoid mating with close kin, such as dispersal or active kin recognition (Widdig et al., 2017), which, as we show here, are effective even in small, isolated populations. Future work could elucidate which factors are most relevant to avoiding close-kin mating or breeding in this important study system. Second, our results show that the Cayo rhesus macaque population falls within the range of natural rhesus macaque populations, both in the prevalence of close-kin breeding and in genomic background relatedness. These non-exceptional patterns further highlight this population’s suitability as a study population and provide a base for future studies. If genomic sample sizes become sufficiently large, one could meaningfully correlate behavioural and phenotypic data with ROH-based estimates of inbreeding. Key insights into the consequences of inbreeding and background relatedness, relevant to conservation biology and evolutionary biology more generally, could thus be gleaned. Our study marks a step in that direction by providing a robust ROH-calling pipeline and estimates of ROH of several rhesus macaque populations.

## Data availability

No new genomic data were generated for this study. Aligned sequencing data for the Cayo Santiago samples are available on the NCBI Sequencing Read Archive within project PRJNA1068427 (www.ncbi.nlm.nih.gov/bioproject/1068427) and project PRJNA251548 (www.ncbi.nlm.nih.gov/bioproject/PRJNA251548; Freudiger et al., 2025). Raw sequencing data for the Chinese samples are available in the NCBI Sequence Read Archive within project PRJNA345528 (www.ncbi.nlm.nih.gov/bioproject/345528), and the VCF is published on GigaDB within dataset 100484 (https://doi.org/10.5524/100484; Liu et al., 2018). ROH segments inferred in this study are available in **Datasets D3 and D4**.

## Code availability

All code is available on GitHub (https://github.com/flopau76/ROH-Cayo).

## Author contributions

We describe the specific contributions of each author below, following the CRediT (Contributor Roles Taxonomy) framework.

**Florence Pautet:** Conceptualisation; Methodology; Data curation; Formal analysis; Visualisation; Writing – original draft.

**Annika Freudiger:** Data curation, Supervision; Writing - review & editing

**Angelina Ruiz Lambides:** Data curation; Project administration; Resources (field site coordination and oversight); Supervision; Investigation; Writing – review & editing.

**Anja Widdig:** Conceptualisation; Data curation; Supervision; Project administration; Funding acquisition; Writing – original draft.

**Harald Ringbauer:** Conceptualisation; Methodology; Supervision; Funding acquisition; Writing – original draft.

## Acknowledgments

We thank the Caribbean Primate Research Center, especially Melween Martinez, Carlos A. Sariol Curbelo, and all field staff at Cayo Santiago, for collecting genetic and demographic data and for their continued support of our work. We acknowledge the genetic consortium, including Fred Bercovitch, Matt Kessler, John Berard, Michael Krawczak, Peter Nürnberg, and Jörg Schmidtke, for their effort to instigate the Cayo Santiago genetic database. We thank the Cologne Center for Genomics, especially Peter Nürnberg, Kerstin Becker, and Janine Altmüller, for producing the sequencing data, and Vladimir M. Jovanovic, Yilei Huang, Noah Snyder-Mackler, and Katja Nowick for their help with the prior sequencing analysis. The sequencing costs, as well as graduate funding to AF, were provided by the Deutsche Forschungsgemeinschaft (DFG) as part of a sequencing initiative (DFG Grant No. WI 1808/7-1, Project No. 433163659, approved to AW). FP and HR are supported by the European Research Council (ERC, Horizon Europe, grant number 101164941, EPIDEMIC, approved to HR). Cayo Santiago is supported by the Office of Research Infrastructure Programs (ORIP) of the NIH (P40 OD012217). The content of this publication is solely the responsibility of the authors and does not necessarily represent the official views of NIH or ORIP.

## Supplementary Information

### Supplementary Data

**Dataset D1**: Metadata and inbreeding coefficients for the Cayo Santiago individuals

**Dataset D2**: Metadata and inbreeding estimate of the Chinese individuals

**Dataset D3**: Filtered and merged ROH segments in the Cayo Santago individuals

**Dataset D4**: Filtered and merged ROH segments in the Chinese individuals

### Supplementary Tables

**Table S1:** Number of Cayo Santiago individuals in the different kin class categories based on pedigree. This table lists the prevalence of various parent-relatedness classes derived from the pedigree. Parental kin classes are sorted according to their degree of kinship (distance in the pedigree. The numbers are based on the 63 individuals for whom we ran TRACE and for whom both parents and all four grandparents were known. For a kin class of degree *d*, the inbreeding estimate *F_ped* of the offspring is half the level of relatedness *r_ped* of the parents, which can be calculated by *r_ped* = 1/2^d^.

| parental kin class | degree of kinship | number of offsprings |
| --- | --- | --- |
| half 1st-cousins | 4 | 3 |
| half 1st-cousins-once-removed | 5 | 1 |
| half 2nd-cousins | 6 | 4 |
| half 2nd-cousins-once-removed | 7 | 2 |
| half 2nd-cousins-twice-removed | 8 | 7 |
| half 3rd-cousins | 8 | 3 |
| half 3rd-cousins-once-removed | 9 | 16 |
| half 3rd-cousins-twice-removed | 10 | 1 |
| half 4th-cousins | 10 | 6 |
| half 4th-cousins-once-removed | 11 | 5 |
| half 3rd-cousins-thrice-removed | 12 | 1 |
| half 5th-cousins | 12 | 1 |
| unrelated | / | 13 |
| **total** | **/** | **63** |

**Table S2:**
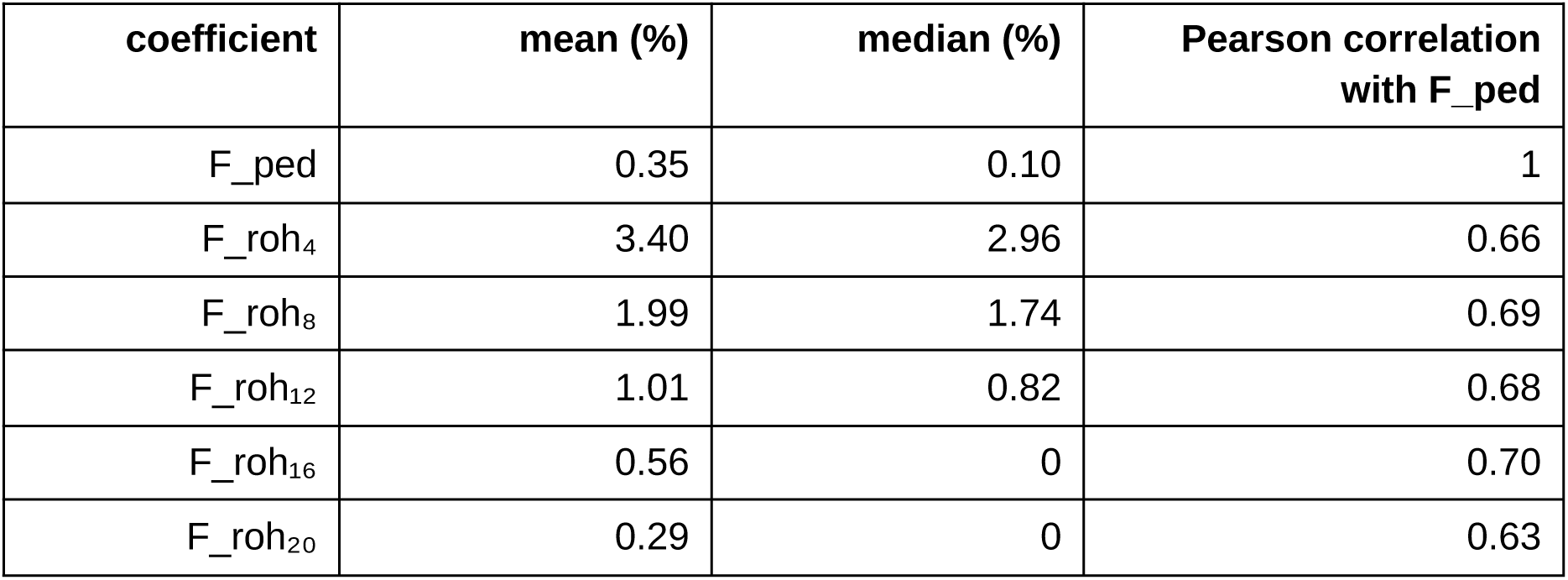
Correlation between F_roh and F_ped on Cayo Santiago. We computed both *F_roh* and *F_ped* for the 63 individuals with complete three-generation pedigrees. For *F_roh*, we used ROH longer than various minimum-length cutoffs.

| coefficient | mean (%) | median (%) | Pearson correlation with F_ped |
| --- | --- | --- | --- |
| F_ped | 0.35 | 0.10 | 1 |
| F_roh<sub>4</sub> | 3.40 | 2.96 | 0.66 |
| F_roh<sub>8</sub> | 1.99 | 1.74 | 0.69 |
| F_roh<sub>12</sub> | 1.01 | 0.82 | 0.68 |
| F_roh<sub>16</sub> | 0.56 | 0 | 0.70 |
| F_roh<sub>20</sub> | 0.29 | 0 | 0.63 |

### Supplementary Figures

**Figure S1:**
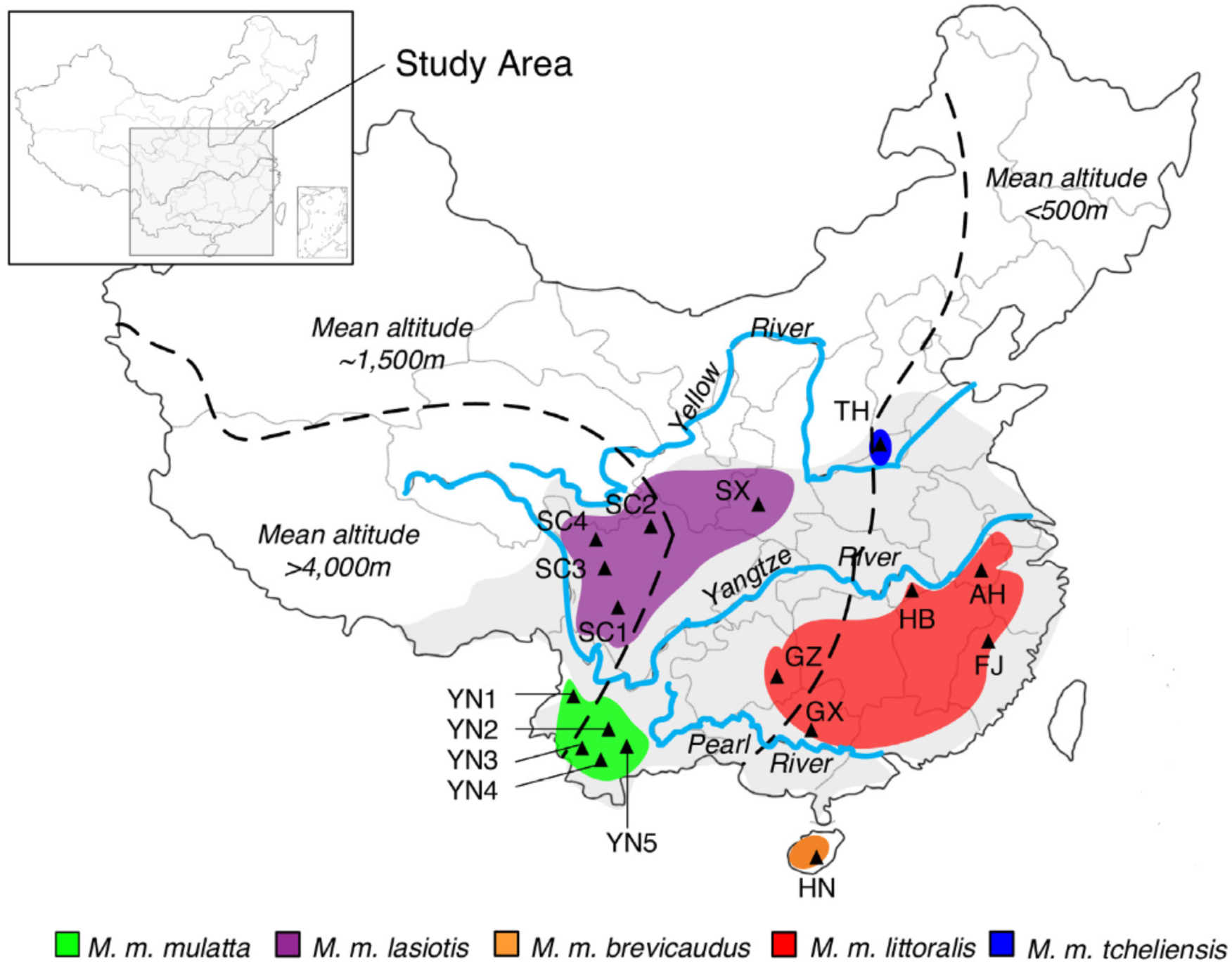
Geographic distribution of five Chinese rhesus macaque populations analysed in this study. The native range of the rhesus macaque stretches across present-day China (shown in grey). The map depicts the 17 sampling sites within their population areas. Figure adapted from (Liu et al., 2018) published in *GigaScience*, licensed under CC BY 4.0.

**Figure S2:**
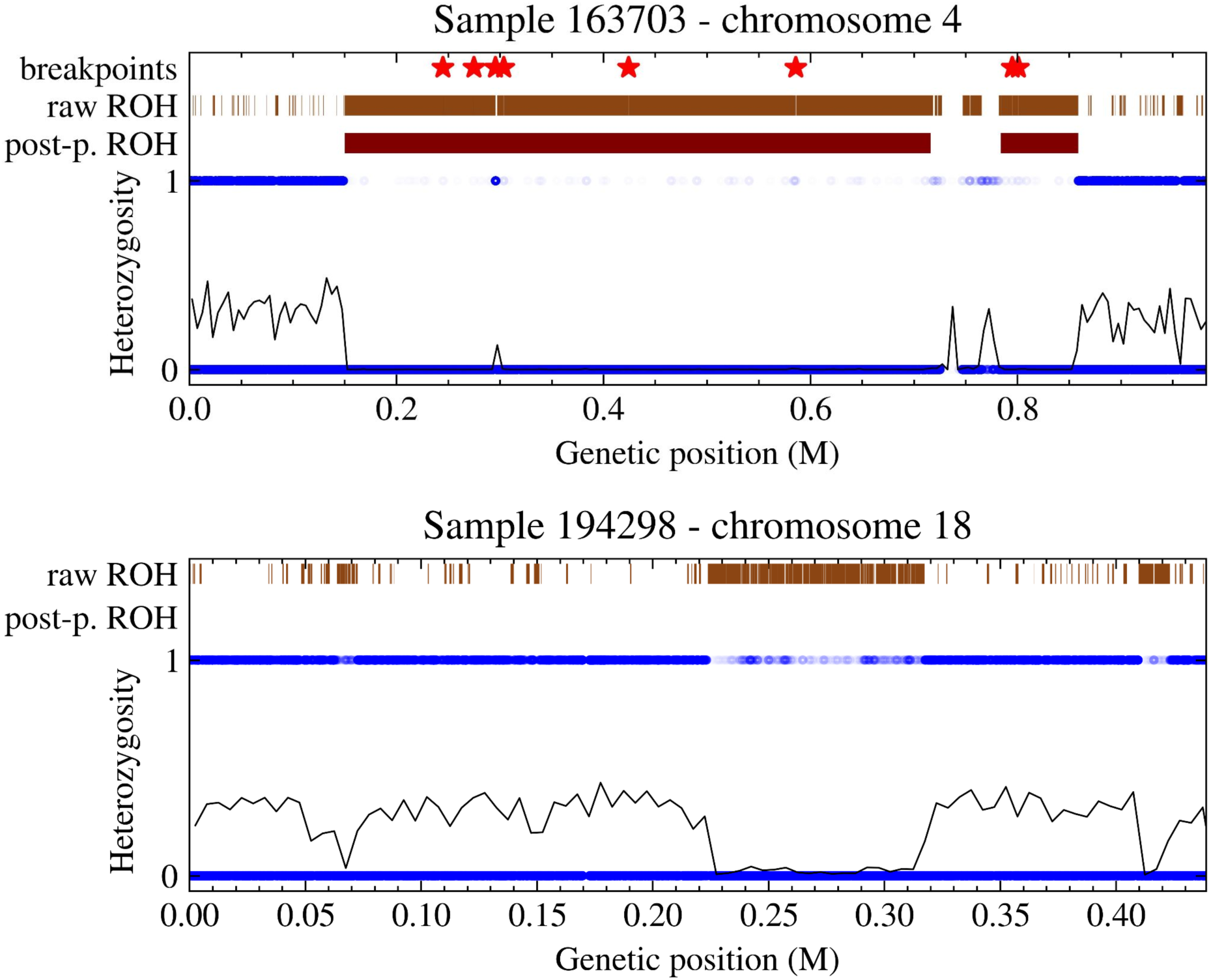
Comparison of inferred ROH before and after post-processing. We compare inferred ROH with heterozygosity at individual markers: at 10,000 randomly selected SNPs (depicted by blue dots) and averaged values over sliding windows of width 0.5 cM (shown as black lines). The inferred ROH are depicted above for both the raw *BCFtools* input and for the post-processed segments. **a:** A single long ROH is clearly visible from a continuous stretch of exceptionally low density of heterozygous SNPs. However, the inferred ROH is divided into 7 inferred ROH, separated by gaps < 1 cM and marked by stars. Those fragments are successfully merged during post-processing. **b**: This genome (Sample 194298) was excluded due to likely contamination. A ROH fragment is evidently visible, stretching between 0.25M and 0.30M, but is not called correctly by our algorithm due to an irregularly elevated number of heterozygous SNPs. Such false heterozygous SNPs may result from DNA contamination or high genotyping error rates.

**Figure S3:**
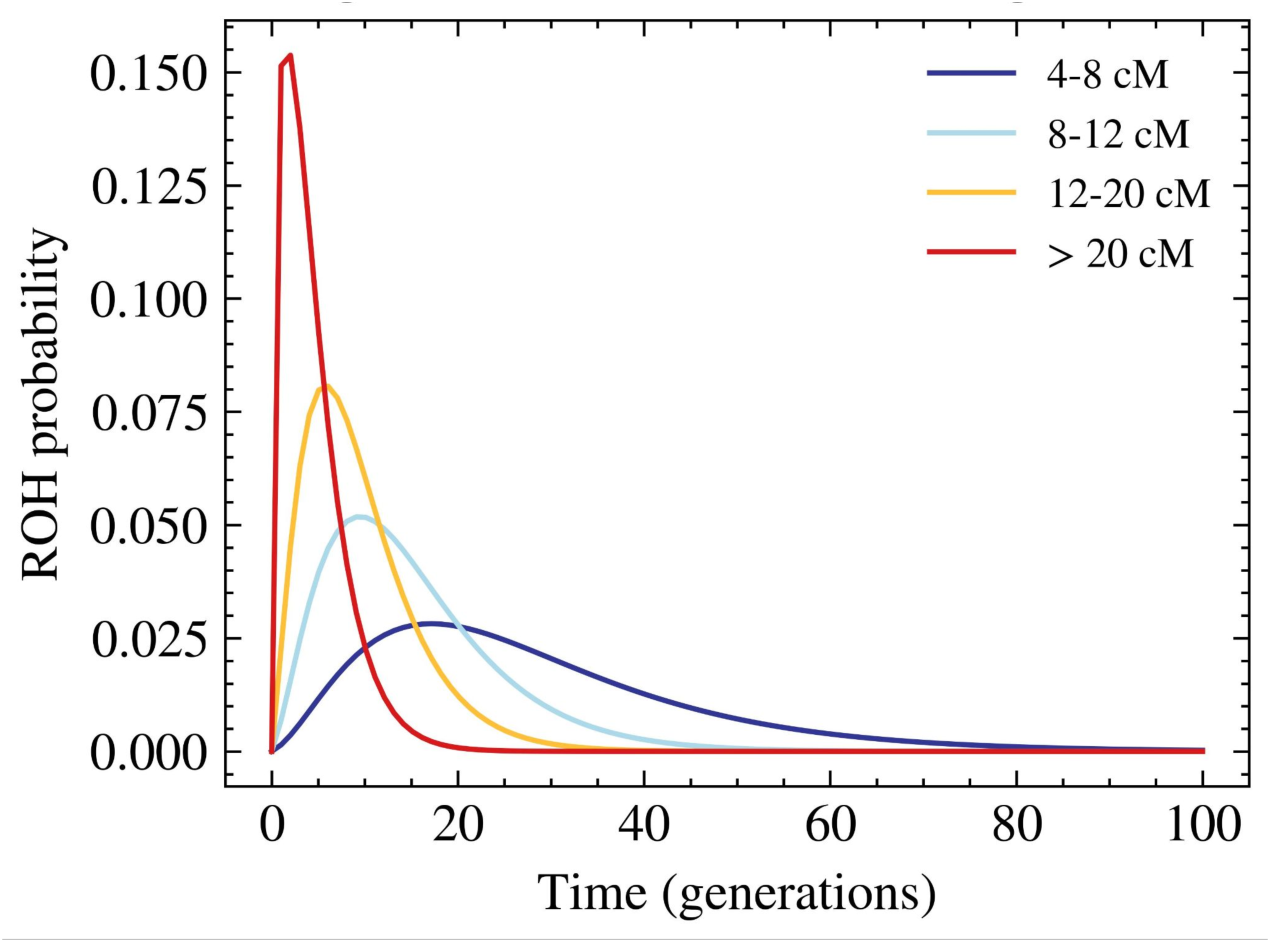
Timescales of coalescence events creating ROH under a constant population size. We plot the probability density of segments in different bins originating at a given number of generations ago. It was computed for a panmictic population of constant size N_e_ = 400, using the *rheMac10* genetic map of Freudiger et al. (2025). We note that the exact population size N_e_ and chromosome lengths have only a very small effect (when N_e_ >> 1).

**Figure S4:**
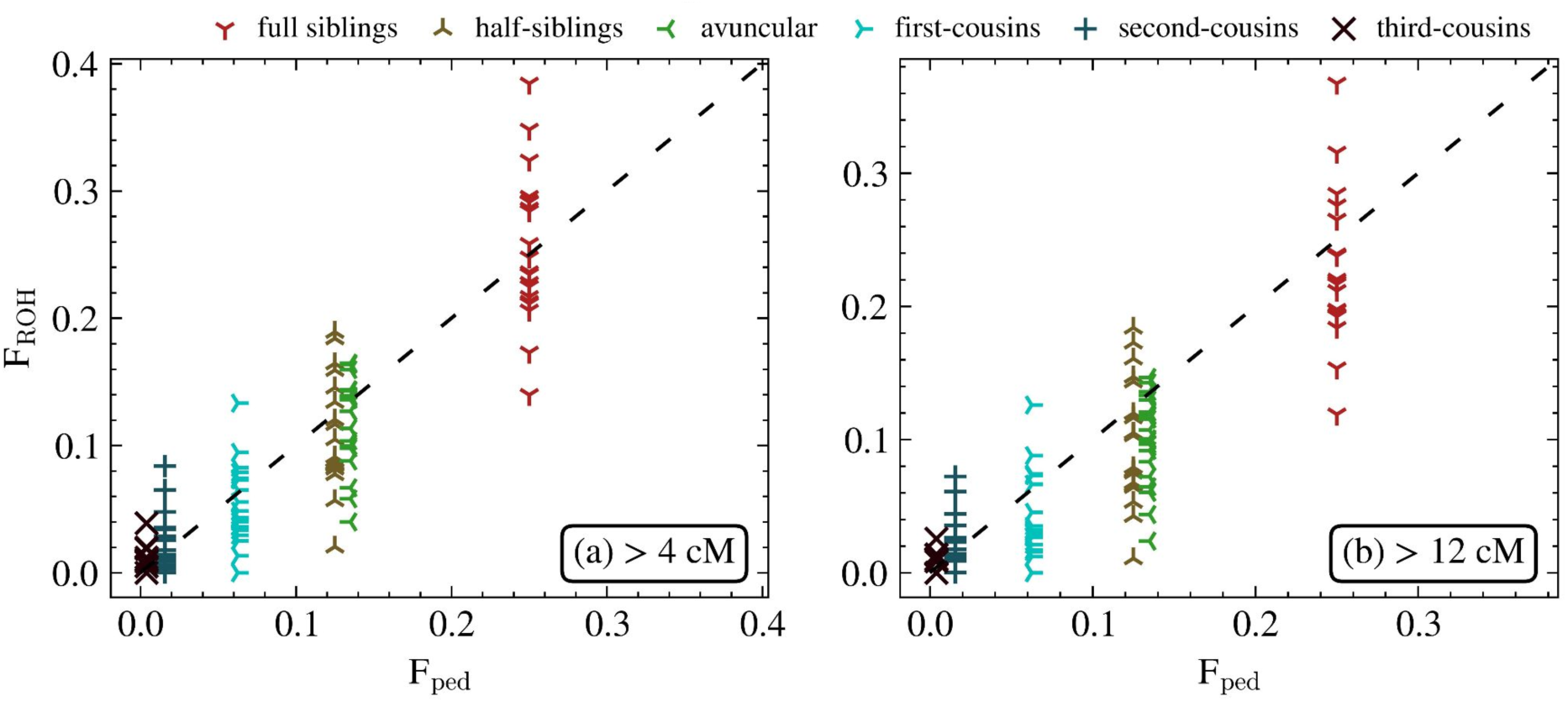
Comparison of inbreeding coefficients F_ped and F_roh in simulated data. Each dot represents an individual simulated using a specific pedigree with *pedsim*, where the kinlabel denotes the degree of relatedness of the parent. F_roh was computed using all segments **(a)** > 4 cM and **(b)** > 12 cM. The dashed line represents the agreement F_ped = F_roh. For visualisation, offspring of avuncular pairs were shifted slightly to the right, even though they have the same F_ped = 0.125 as offspring of half-siblings.

**Figure S5:**
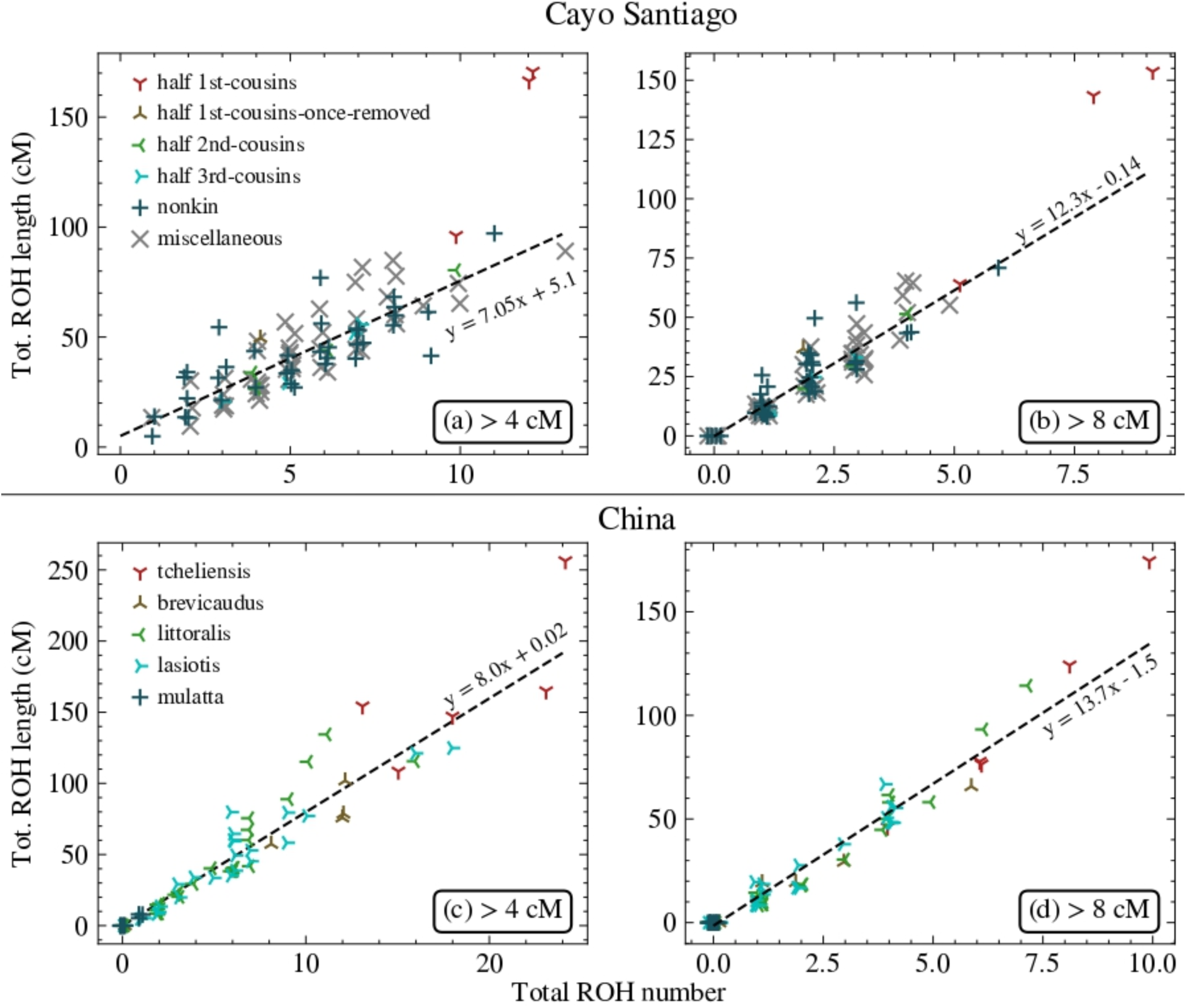
Distribution of the number and size of ROH segments in Cayo Santiago and Chinese rhesus macaques. **(a)** and **(c)** show data for ROH segments > 4cM and **(b)** and **(d)** for ROH segments > 12 cM, for the Cayo Santiago **(a)** and **(b)** and the Chinese samples **(c)** and **(d)**. Each dot represents one individual. The dashed line corresponds to the least-squares linear regression, excluding highly inbred samples (more than 5% of the genome in ROH > 16 cM long). For visualisation, random horizontal jitter was added to **the** points.

## Notes

https://github.com/flopau76/ROH-Cayo

### Competing Interest Statement

The authors have declared no competing interest.

### Summary of Updates

Fixed numeric values in Table S2 and made minor orthographic corrections.

